# Impact of gradient scheme and shimming on out-of-voxel echo artifacts in edited MRS

**DOI:** 10.1101/2022.07.10.499491

**Authors:** Yulu Song, Helge J. Zöllner, Steve C.N. Hui, Kathleen Hupfeld, Georg Oeltzschner, Richard A.E. Edden

## Abstract

Out-of-voxel (OOV) signals are common spurious echo artifacts in magnetic resonance spectroscopy (MRS). These signals often manifest in the spectrum as very strong ‘ripples’ which interfere with spectral quantification by overlapping with targeted metabolite resonances. Dephasing Optimization Through Coherence Order Pathway Selection (DOTCOPS) gradient schemes are algorithmically optimized to suppress all potential alternative coherence transfer pathways (CTPs), and should suppress unwanted OOV echoes. In addition, second-order shimming uses non-linear gradient fields to maximize field homogeneity inside the voxel, which unfortunately increases the diversity of local gradient fields outside of the voxel. Given that strong local spatial B_0_ gradients can refocus unintended CTPs, it is possible that OOVs are less prevalent when only linear first-order shimming is applied. Here we compare the size of unwanted OOV signals in Hadamard-edited (HERMES) data acquired with either a local gradient scheme (which we refer to here as “Shared”) or DOTCOPS, and with first or second-order shimming. We collected data from 15 healthy volunteers in two brain regions (voxel size 30×26×26mm^3^) from which it is challenging to acquire MRS data: medial prefrontal cortex (MPFC) and left temporal cortex (LTC). Characteristic OOV echoes were seen in both GABA- and GSH-edited spectra for both brain regions, gradient schemes, and shimming approaches. A linear mixed-effect (LME) model revealed a statistically significant difference in the average residual based on gradient scheme in both GABA-(p < 0.001) and GSH-edited (p < 0.001) spectra; that is, the DOTCOPS gradient scheme resulted in smaller OOV artifacts compared with the Shared scheme. There were no significant differences in OOV artifacts associated with shimming method. Thus, these results suggest that the DOTCOPS gradient scheme for *J*-difference-edited PRESS acquisitions yields spectra with smaller OOV echo artifacts than the Shared gradient scheme implemented in a widely disseminated editing sequence.

## 1. Introduction

*J*-difference-edited magnetic resonance spectroscopy (MRS) is a noninvasive quantitative method to detect signals from endogenous tissue metabolites, particularly overlapped signals from less concentrated metabolites^1^. As a difference method that subtracts large signals to reveal small signals of interest, edited MRS requires high-quality data^2^. One common artifact seen in single-voxel difference spectra is out-of-voxel (OOV) water echoes, or ‘ghosts’^3^. Water signals from outside the voxel, which are not sufficiently removed by pre-saturation water suppression or the coherence transfer pathway (CTP) selection^4^, can take unintended pathways through the sequence from excitation to detection. These often result in echoes that are not synchronized with the beginning of the acquisition window, and thus give rise to broad signals in the spectrum with strong first-order phase ‘ripple’. As these signals arise from outside the shimmed voxel, they can occur at any chemical shift, and interfere with quantification of within-voxel signals.

MRS sequences employ CTP selection in order to suppress unintended signals and retain only the signals of interest. Both phase cycling and dephasing gradient (or ‘crusher gradient’) pulse schemes can achieve CTP selection^5^; however, in the context of in vivo MRS^6^ (where the ‘sample’ is not static), gradient pulse schemes are particularly important. The PRESS pulse sequence shown in Figure 1, has a gradient scheme that is designed to select the desired CTP [-1, +1, -1] as shown in black, and suppress other pathways. One example of a CTP that could result in unwanted OOV echoes is also shown [+1, +1, -1] in red. Although the gradient scheme shown should efficiently suppress this CTP, B_0_ gradients outside of the voxel (whether they arise from local anatomy or second-order shim fields^7^) can potentially refocus such pathways. The effectiveness of a given gradient scheme for suppressing OOV echoes can be improved by changing the order, amplitude and polarity of slice-selective gradients^8^. Recently, Dephasing Optimization Through Coherence Order Pathway Selection (DOTCOPS)^9^ was developed to optimize gradient schemes algorithmically, so as to suppress all potential alternative CTPs. DOTCOPS-optimized gradient schemes have been proposed for most commonly used MRS sequences, including spectral-editing (MEGA) sequences with PRESS and sLASER localization^9^.

**Figure 1.**
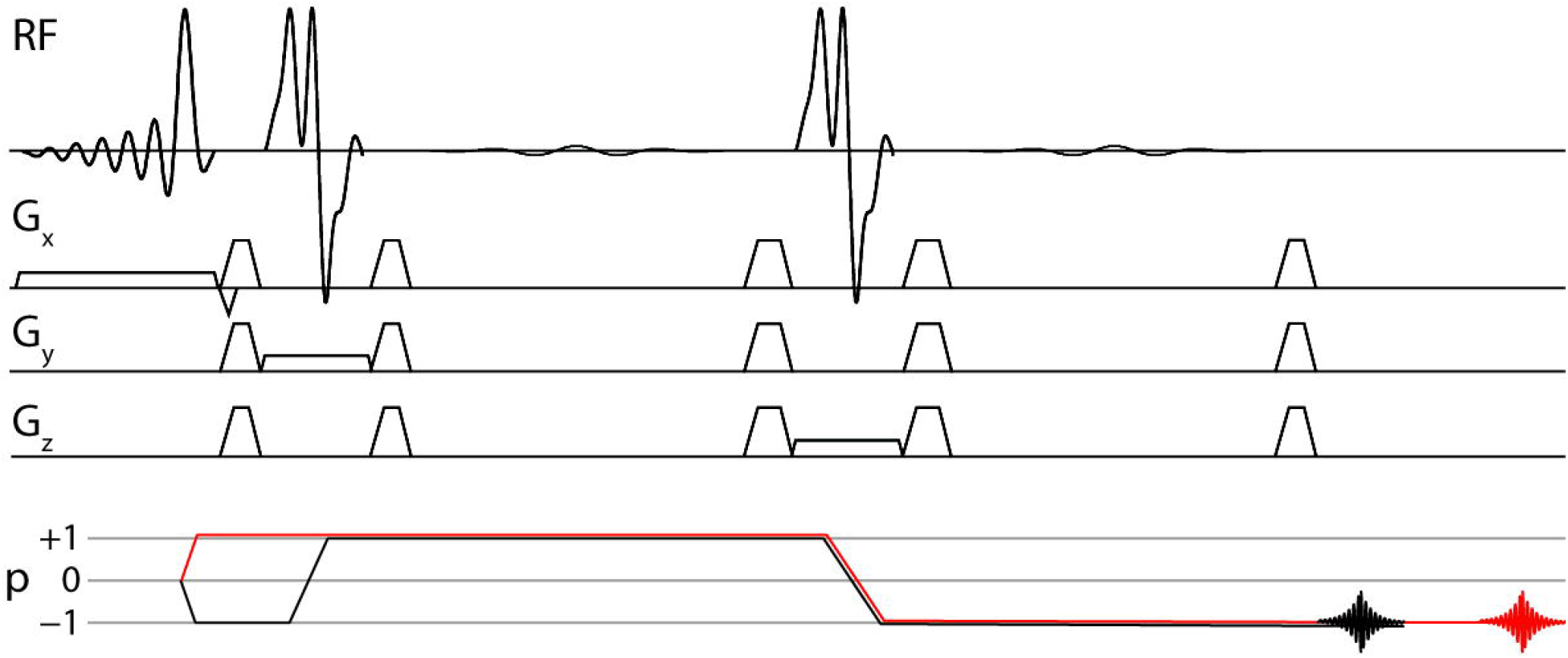
PRESS pulses sequence and coherence transfer pathway (CTP) diagram. The intended CTP is shown in black, and one example of an undesirable coherence pathway is shown in red.

DOTCOPS represents an important advance in gradient scheme design, proposing a theoretical framework for quantitatively selecting one gradient scheme over another, and proposing optimized gradient schemes for a number of common sequences^9^. Given the broad scope of the work, it was not possible to provide in vivo demonstrations of all gradient schemes proposed. A key limitation of the implementation described in the original publication (though not of the DOTCOPS approach in general) is that only CTPs arising from excitation during the single-voxel localization pulses are considered, but not those related to common pre-localization modules such as water suppression and outer-voxel-saturation (OVS). However, as mentioned above, spurious echoes observed in MRS experiments that are refocused into the acquisition window often originate from outside the localized voxel. Any such echo must have been excited before the single-voxel localization module, i.e. by water suppression of OVS pulses. The gradient schemes proposed in ref^9^ do not consider these CTPs, nor their degree of suppression, and might therefore be more susceptible than expected to OOV arising from accidently fold: First, we set out it to test whether the proposed DOTCOPS gradient scheme for MEGA-PRESS experiments is more effective than the gradient scheme that is currently implemented in the HERMES spectral-editing sequences implemented for Philips systems (referred to in the present work as the “Shared” gradient scheme). To do so, we acquired edited MRS data from two challenging brain regions, from which OOV echoes are commonly seen, the medial prefrontal cortex (MPFC) and left temporal cortex (LTC). Second, we also investigated whether the appearance of OOV echoes is affected by the use of first-vs. second-order shim fields, as second-order shim fields could plausibly increase the likelihood of accidentally refocusing unintended CTPs.

## 2. Methods

This manuscript compares the effect of two gradient schemes and two shimming approaches on OOV artifacts in the context of *J*-difference edited MRS. Specifically, we compare the DOTCOPS MEGA-PRESS scheme^9^ with the Philips ‘Shared’ gradient scheme, which our prior work^10–14^ has generally used for Philips implementation of edited MRS with PRESS localization, and which has been widely shared. We developed the Shared gradient scheme: in the Philips product PRESS sequence, each slice-selective refocusing pulse is surrounded by a pair of gradients on each of the three axes (labeled *x, y*, and *z* for convenience, but rotating with the voxel). In order to develop this into a sequence for *J*-difference editing, we moved one gradient from before each editing pulse to after it, resulting in the scheme shown at the bottom in Figure 2A.

**Figure 2.**
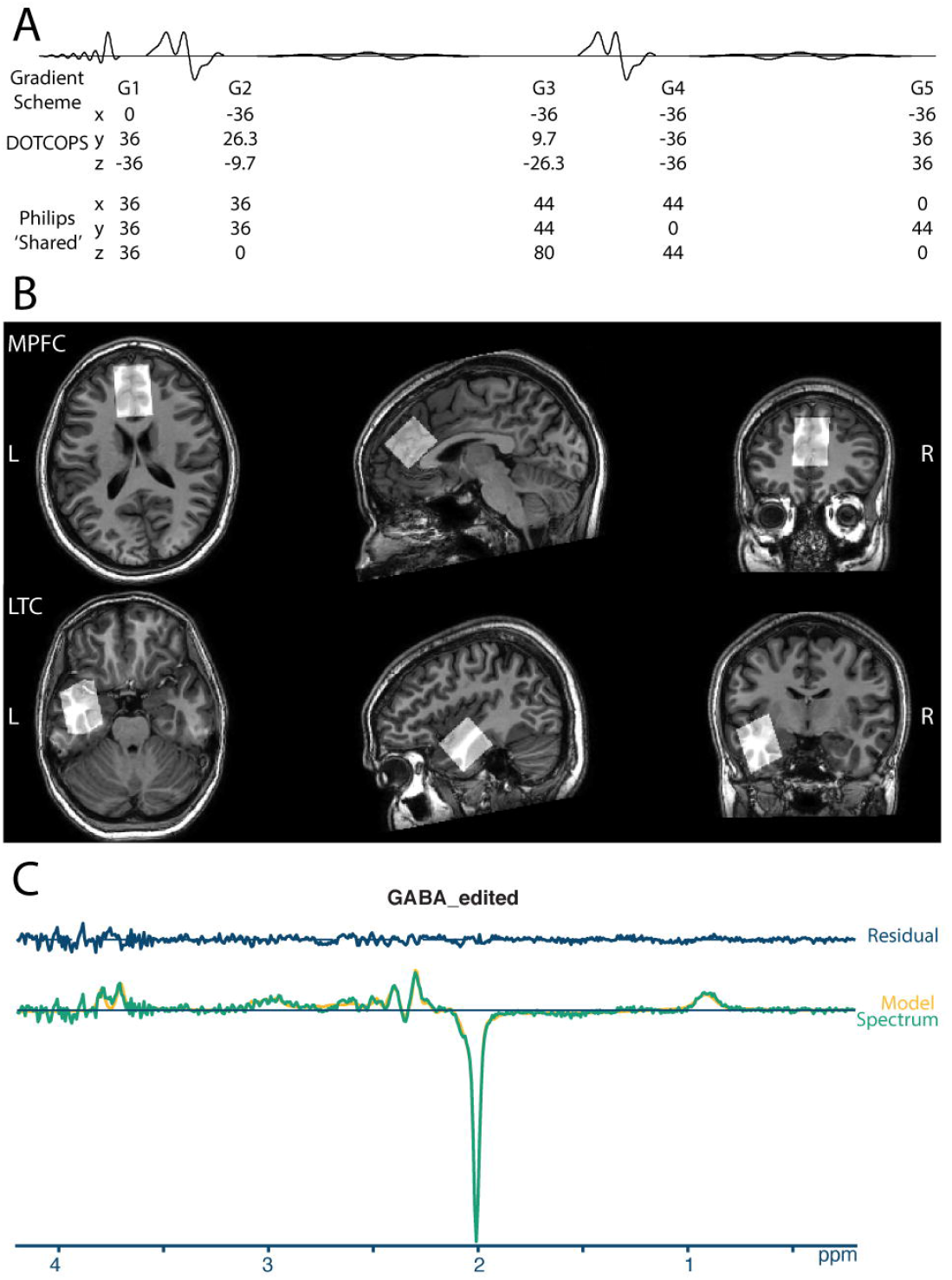
A. DOTCOPS and Philips “Shared” Gradient schemes. Values are gradient areas in units of mT.ms/m. B. In vivo HERMES spectra were acquired from medial prefrontal (MPFC) and left temporal cortex (LTC) voxels (30 × 26 × 26 mm^3^), shown here for a single exemplar subject. C. GABA-edited difference spectrum and Osprey model for DOTCOPS data from a single exemplar subject. This spectrum shows several small OOV echoes between 3.5 and 4 ppm. Gold line: Model. Green line: Spectrum. Blue line: Model residual (i.e., the difference in Spectrum-Model).

The use of a gradient scheme to suppress unwanted OOV echoes has one potential weakness. B_0_ inhomogeneity outside the voxel of interest, which can be thought of as a spatially varying field gradient, might interact with the gradient scheme to refocus unintended CTPs, generating OOV echoes. Second-order shimming uses non-linear gradient fields to maximize field homogeneity inside the voxel, increasing the diversity of local gradient fields outside of the voxel. Here we hypothesis that OOV echoes might be less prevalent when only first-order shimming is employed, and more prevalent with second-order shimming.

In order to test the success of each gradient scheme for the specific task of suppressing OOV echo signals in J-difference-edited spectra, we use the HERMES experiment^15^. This has the advantage of yielding two difference-edited spectra (i.e., both GABA- and GSH-edited spectra) in a single acquisition. Since the HERMES acquisition includes multiple frequency-selective editing pulses at different frequencies, it is possible that different sets of potential unwanted echoes are refocused.

### 2.1 Participants

Fifteen healthy volunteers (4M/11F, average age: 23.8 ± 2.8 years old) provided their written informed consent to participate. All data were acquired with approval from the local Institutional Review Board.

### 2.2 Data acquisition

HERMES data were acquired on a Philips 3T Elition RX scanner (Philips Healthcare, Best, the Netherlands) equipped with a 32-channel head coil. The scanning protocol included a 1 mm^3^ isotropic T_1_-weighted image (MPRAGE) scan for voxel placement (TR 8.0 ms, TE 3.7 ms, flip angle 8°). Next, two 30 × 26 × 26 mm^3^ (AP x LR x HF) voxels were placed, in the medial prefrontal (MPFC) and left temporal cortex (LTC) regions as shown in Figure 2B. These two regions are of substantial interest in studies of psychiatric and neurogenerative disorders^16–18^, and are particularly challenging for MRS^7^ due to their close proximity to tissue-to-air interfaces and resulting discontinuity in magnetic susceptibility. MPFC data were acquired in 15 subjects, and LTC data in 10 subjects. For each voxel location, four separate HERMES experiments were performed: using either the DOTCOPS or Shared gradient scheme; and with the Philips implementation of pencil-beam shimming either to the first (linear gradients) or second (quadratic gradients) order. For each experiment, a total of 160 transients were acquired in 5min 36s, using the following parameters: TR 2000 ms, TE = 80 ms; 14 ms editing pulses applied in a Hadamard-encoded scheme at 1.9 ppm (GABA-ON) and 4.56 ppm (GSH-ON); 1024 datapoints; 2 kHz spectral width; readout duration is 512 ms; and MOIST water suppression^19^.

### 2.3 Data analysis

Data were analyzed in Osprey^20^ using consensus recommended pre-processing^21^ and linear-combination modeling. We have recently shown that linear-combination modeling is superior to simpler modeling approaches for HERMES data^13^. In brief, the frequency-and-phase alignment of FIDs was performed using robust spectral registration^22^. A Hankel singular value decomposition (HSVD) filter^23^ was applied to remove residual water signals and to reduce baseline roll. Spectra were modeled between 0.2 and 4.2 ppm. Twenty-five metabolite basis functions were included in the model (Asc, Asp, Cr, GABA, GPC, GSH, Gln, Glu, Ins, Lac, NAA, NAAG, PCh, PCr, PE, sI, Tau) with 8 MM basis functions in the sum spectrum (MM_0.94_, MM_1.22_, MM_1.43_, MM_1.70_, MM_2.05_, Lip09, Lip13, Lip20) parameterized as Gaussians. For the GSH-edited difference spectra, co-edited macromolecule (MM) peaks defined at 1.2 and 1.4 ppm. Similarly, two MM basis functions were added at 0.93 and 3 ppm for the GABA-edited difference spectrum. Amplitude-ratio soft constraints are imposed on the MM and lipid amplitudes, and NAAG/NAA amplitudes, as defined in the LCModel manual^24^. The Osprey’s default baseline knot spacing of 0.4 ppm was used for the cubic spline baseline. The co-edited MM3co amplitude had a 1-to-1 soft constraint to the GABA amplitude (‘1to1GABAsoft’ setting in Osprey) which has been shown to improve modeling performance^13,21,25,26^. The water-reference data were quantified with a simulated water basis function in the frequency domain with a six-parameter model (amplitude, zero- and first-order phase, Gaussian and Lorentzian line broadening, and frequency shift). A brief summary of experimental methods following the minimum reporting standards paper in MRS^27^ is added in Supplementary Material 1.

### 2.4 Quantification of OOV artifacts

In order to compare gradient schemes and shimming effects, a metric of artifact size is required. Since OOV echoes are the largest contributor of unmodeled signal where present, we used the minimum to maximum amplitude of the linear-combination-modeling residual normalized by the noise as a quantitative metric of OOV signal. Figure 2C shows the model and model-residual for one spectrum with small, but clearly identifiable OOV echoes.

### 2.5 Statistical analysis

A robust regression linear mixed-effects (LME) model was performed to test whether residual estimates from OOV echoes differed between the DOTCOPS and Shared gradient schemes, or between first- and second-order shimming. In this case, where the occurrence of occasional ‘very-bad’ outliers is of specific interest, ANOVA analysis is not appropriate. LME models use the linear regression paradigm^28^ to parsimoniously describe data that are collected and summarized in groups^29^. Robust regression^30,31^ is an alternative to least squares regression when data is contaminated with outliers or influential observations. Therefore, robust regression is appropriate for this dataset. We considered statistical significance as p < 0.05.

## 3. Results

All spectra were successfully analyzed using Osprey. Edited difference spectra, overlaid for all subjects, are shown in Figure 3. Characteristic OOV echoes can be seen in both GABA- and GSH-edited spectra for both regions, gradient schemes, and shimming approaches. Note that these spectra are presented without line-broadening which can dampen the amplitude of OOV echoes that occur late in the FID. The relative amplitude of the model residuals, which we interpret as a measure of OOV echo amplitude, are plotted in Figure 4. Note that the open white circle in each case is the group mean.

**Figure 3.**
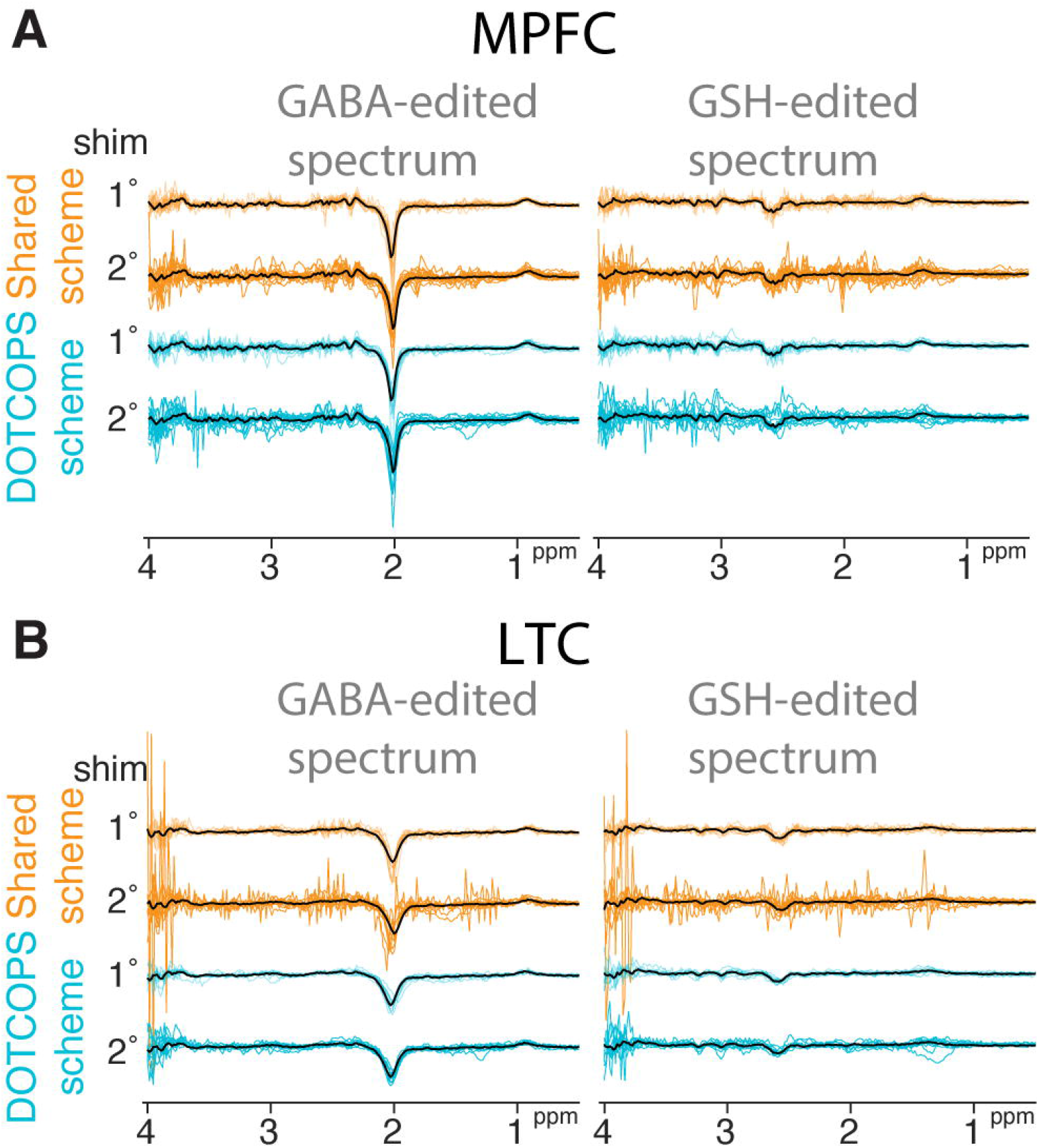
GABA- and GSH-edited difference spectra overlaid from all subjects, for two regions: **A** medial prefrontal (MPFC) and **B** left temporal cortex (LTC). Spectra acquired with the Shared and DOTCOPS gradient schemes are depicted in gold and cyan, respectively. Mean spectra calculated across subjects are depicted in black. Data acquired with shimming to the first- or second-order are labeled 1° and 2°, respectively.

**Figure 4.**
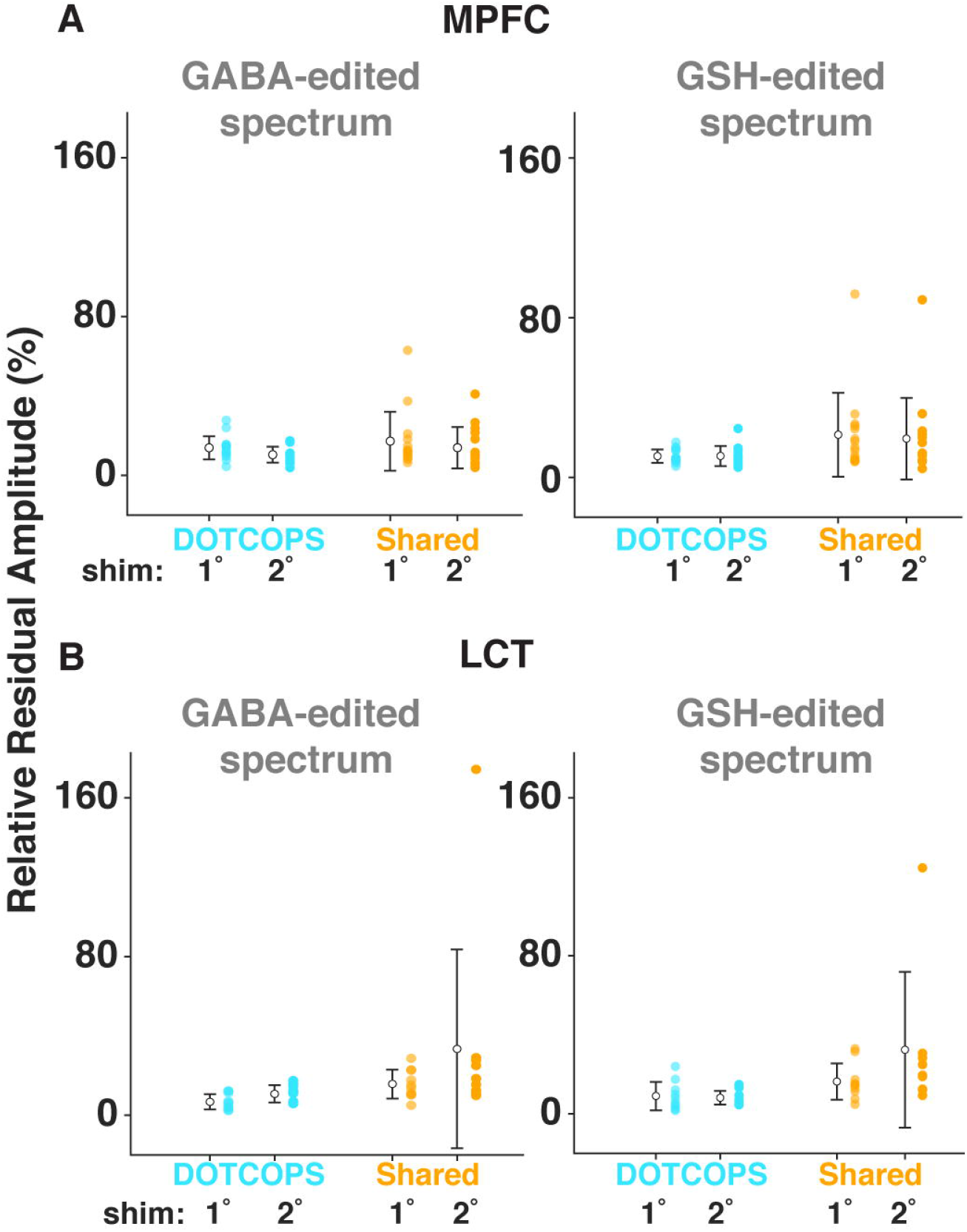
Ball and whisker plots of the model residuals of the two gradient schemes, shimming orders, and brain regions for both GABA- and GSH-edited spectra. (A) medial prefrontal cortex (MPFC) voxel; (B) left temporal cortex (LTC) voxel. Data acquired with the DOTCOPS and Shared gradient schemes are depicted in cyan and gold, respectively. Data acquired with shimming to the first- or second-order are labeled 1° and 2°, respectively. Open white circles indicate group-mean values. Whiskers represent mean ± SD.

The LME model revealed a statistically significant difference in average residual for gradient scheme, for both GABA (p < 0.001) and GSH (p < 0.001). That is, the DOTCOPS scheme yielded smaller OOV artifacts compared with the Shared scheme. The interaction of region*gradient scheme was significant for GABA (p < 0.01), but not for GSH (p >0.05), as shown in Table 1. In the GABA-edited spectra, region also has a significant effect, in the direction that LTC region has worse artifacts than MPFC. No significant differences were found between the shimming schemes.

**Table 1.** LME results. Separate models were run for the GABA- and GSH-edited spectra. In each case, the factor *gradient* compared the DOTCOPS vs. Shared schemes, the factor *region* compared the MPFC vs. LTC voxels, and the factor *shim* compared spectra acquired after first- or second-order shimming.

|  |  | P value |
| --- | --- | --- |
| GABA residual | gradient | <0.001 |
|  | region | 0.029 |
|  | shim | 0.992 |
|  | gradient * region | 0.007 |
| GSH residual | gradient | <0.001 |
|  | region | 0.162 |
|  | shim | 0.755 |
|  | gradient * region | 0.060 |

## 4. Discussion

Obtaining high-quality edited MRS data is critical, especially when trying to measure metabolites present at low-concentration in the human brain^1^. OOV echoes are a common artifact seen in edited spectra that should ideally be suppressed by CTP selection. Pulsed field gradients^32^ achieve CTP selection within each transient, rather than by subtracting unwanted CTPs across a phase cycle. This approach is particularly preferable for edited MRS, where frequency and phase correction (FPC) is a necessary pre-processing step, because there is a conflict between the assumptions of phase cycling and most FPC algorithms^33^. The disadvantage of using gradients for CTP is that they must be accommodated during the echo time - it is therefore very important to develop gradient pulse schemes that efficiently suppress unwanted coherences. The DOTCOPS algorithm approaches this question in a systematic fashion, leading to the proposed gradient scheme for edited PRESS experiments. Even though the CTPs of OOV echoes excited by water suppression pulses were not fully considered in the DOTCOPS optimization, the DOTCOPS gradient scheme outperformed the Shared gradient scheme that our Philips-based sequences have used to date.

Single-voxel MRS is not equally easy throughout the brain. Tissue-bone-air interfaces lead to complex local distortions of the B_0_ field, that are imperfectly addressed by shimming, and impact spectral quality by increasing linewidths and making OOV echoes more likely^7^. The interaction between shimming and OOV echoes is complex. That is, shimming routines typically optimize B_0_ field homogeneity *within* the voxel of interest, without consideration for OOV B_0_ field homogeneity. Inhomogeneous local B_0_, or local field gradients, can partially cancel the effect of CTP gradients, and can even lead to refocusing of unwanted echoes within the acquisition window^3^. In this study, we tested whether OOV echoes were more likely to occur when second-order shim terms were used in addition to linear shims, but we found no differences in OOV echoes based on shimming approach.

In addition to efforts to minimize OOV echoes during acquisition, there are several postprocessing methods for reducing these spurious echoes. Some OOV echoes can be refocused hundreds of milliseconds into the acquisition window, when the FID has decayed away; such signals can be removed by time-domain truncation^34^. Most OOV echoes have a refocusing point that lies after, but during, the intended echo – these signals can be relatively reduced by apodization^34^ (e.g., exponential line broadening). Estimation and removal of artifacts using sensitivity encoding (ERASE)^35^ can remove OOV echoes if their spatial origin is known, using phased-array receive-coil sensitivity profiles to separate OOV echo signals from metabolite signals arising from within the voxel^35^. Given that OOV signals can often be identified by eye in both the time- and frequency-domain, there is reason to expect that deep learning techniques will be able to identify and remove spurious echoes. To date, some progress has been made in the removal of OOV signals without human intervention^36^. Further development of such post-processing methods will continue to be valuable, working in parallel to efforts to optimize acquisition methods.

Systematic studies of OOV echoes are challenging. B_0_ field distortions change from subject-to-subject (due e.g., to different sinus geometry) and even from scan-to-scan within subjects (e.g., changing with head orientation). The quantification of artifacts is a further challenge; there is a concern that relying on modeling residual might mix in effects other than OOV echoes. Moreover, statistical testing on artifacts is difficult, as absolute-mode quantification of artifacts yields asymmetric distributions, and positive outliers are the focus of the investigation (and should not be down-weighted or omitted). The cohort size used here (15 in MPFC; 10 in temporal) may not be sufficient to study such variable within- and between-subject effects; thus, further work in larger samples is warranted.

There are several additional limitations to the generalizability of this work. Although the brain regions chosen are two of the most challenging in the brain, and thus most important for addressing OOV echoes, they each have idiosyncratic local field distortions, and findings may differ in other regions. The scope of this investigation includes only two gradient schemes, and the two schemes are ‘balanced’ within that comparison – both in terms of the maximum gradient area and total gradient area, the DOTCOPS scheme is perhaps disadvantaged in the comparison. There is likely to be further benefit to increasing the scope of the DOTCOPS optimization to include pathways excited by, and gradients contained within, water suppression and spatial pre-saturation sequence modules. Notwithstanding these disadvantages, the DOTCOPS gradient scheme for *J*-difference-edited PRESS acquisitions did successfully reduce OOV echo artifacts compared to the gradient scheme used in our widely disseminated Shared editing sequence. Future implementation of the DOTCOPS gradient scheme may improve quantification of metabolites when challenging brain regions are sampled.

## 5. Conclusion

This study compares two gradient schemes and shimming approaches applied to reduce OOV artifacts in edited MRS experiments. The DOTCOPS gradient scheme yielded significantly smaller OOV echo artifacts compared with the commonly-used Shared gradient scheme. There were no differences in OOV artifacts between first- and second-order shimming. Although the current sample size is modest, and the parameters sampled were not exhaustive, these results motivate switching to the DOTCOPS scheme for future edited PRESS experiments.

## Abbreviations

OOV: Out-of-voxel
DOTCOPS: Dephasing Optimization Through Coherence Order Pathway Selection
CTP: coherence transfer pathway
HERMES: Hadamard Encoding and Reconstruction of MEGA-Edited Spectroscopy
OVS: Outer voxel saturation
MPFC: medial pre-frontal cortex
LTC: left temporal cortex
LME: linear mixed-effect
MEGA-PRESS: Mescher–Garwood Point Resolved Spectroscopy
MPRAGE: Magnetization Prepared - RApid Gradient Echo
HSVD: Hankel singular value decomposition
MM: Macromolecule
sI: Scyllo
FPC: frequency and phase correction
ERASE: Estimation and removal of artifacts using sensitivity encoding

## Acknowledgements

This work has been supported by NIH grants P41EB015909, R01EB016089, R01EB023963, K00AG068440, R21AG060245, and R00AG062230.

