## Supplemental Table1 for "Impact of gradient scheme and shimming on out-of-voxel echo artifacts in edited MRS"

| 1.Hardware | |
| --- | --- |
| a. Field strength [T] | 3 T |
| b. Manufacturer | Philips |
| c. Model (software version if available) | R 5.7.1 |
| d. RF coils: nuclei (transmit/receive), number of channels, type, body part | ^1^H, 32 channel, head |
| e. Additional hardware | - |
| 2. Acquisition | |
| a. Pulse sequence | HERMES (Johns Hopkins University Patch) |
| b. Volume of interest (VOI) locations | MPFC, LTC |
| c. Nominal VOI size [mm^3^] | 30 x 26 x 26 mm^3^ |
| d. Repetition time (TR), echo time (TE)[ms] | TR 2000 ms, TE 80 ms |
| e. Total number of averages per spectrum  i. Number of averaged specra per subspectrum | 160 total averages with 40 averages per subspectrum |
| f. Additional sequence parameters  i. Editing pulse parameters | F1: 2000 Hz, 1024 points  GABA at 1.9 ppm, GSH at 4.56 ppm |
| g. Water suppression method | MOIST |
| h. Shimming method, reference peak, and threshold of acceptance of shim chosen | 1^st^ and 2^nd^ shimming pencil beam, water |
| i. Trigger or motion correction | No trigger or active motion correction |
| 3. Data analysis methods and outputs | |
| a. Analysis software | Osprey 1.1.0 |
| b. Processing steps deviating from Osprey | Final alignment of the averaged sub-spectra by  minimizing the choline (not water) peak; co-edited MMs at 3 ppm were modelled using the  “1to1GABA” model |
| c. Output measure | tCr, rawWaterScaled, CSFWaterScaled, TissCorrWaterScaled |
| d. Quantification references and assumptions, fitting model assumptions | Asc, Asp, Cr, GABA, GPC, GSH, Gln, Glu, Ins, Lac, NAA, NAAG, PCh, PCr, PE, sI, Tau, 8 MM basis functions in the sum spectrum (MM_0.94_, MM_1.22_, MM_1.43_, MM_1.70_, MM_2.05_, Lip09, Lip13, Lip20) Fitting method: Osprey baseline knot spacing 0.4 ppm |
| 4. Data quality | |
| HERMES, MPFC | |
| a. SNR (NAA), linewidth (NAA) [Hz, OFF spectra] | SNR: 64 ± 28, linewidth 20.84 ± 23.48 Hz |
| c. Quality measures of postporcessing model fitting (Mean Relative Amplitude Residual (Residual/Noise))  sum  diff1  diff2 | 14.92 ± 12.37  13.87 ± 9.76  15.54 ± 15.44 |
| d. Mean spectrum created with OspreyOverview | Figure 3 |
| HERMES, LTC | |
| a. SNR (NAA), linewidth (NAA) [Hz, OFF spectra] | SNR: 40 ± 11, linewidth 28.64 ± 47.43 Hz |
| c. Quality measures of postporcessing model fitting (Mean Relative Amplitude Residual (Residual/Noise))  sum  diff1  diff2 | 20.02 ± 19.44  16.67 ± 26.57  16.51 ± 22.14 |
| d. Mean spectrum created with OspreyOverview | Figure 3 |

Supplementary Material 1: Summary following minimum reporting standards in MRS^27^ generated in Osprey See Lin et al. 'Minimum Reporting Standards for in vivo Magnetic Resonance Spectroscopy (MRSinMRS): Experts' consensus recommendations. NMR in Biomedicine. 2021;e4484. [doi.org/10.1002/nbm.4448](https://doi.org/10.1002/nbm.4448)
